# Integrating Millions of Years of Evolutionary Information into Protein Structure Models for Function Prediction

**DOI:** 10.1101/2025.11.11.687944

**Authors:** Runze Ma, Chengxin He, Zhongyue Zhang, Huiru Zheng, Lei Duan

## Abstract

**Background:** Understanding life processes relies on accurate protein function prediction, fundamentally requiring the integration of evolutionary information encoded in sequences with spatial characteristics from 3D structures. Existing approaches often face limitations, however, by either over-relying on sequence, using simplified structural representations instead of fine-grained spatial details, or failing to capture the synergistic relationship between sequence and structure, compounded by challenges in acquiring annotated data.

**Results:** To address these issues, we propose a novel contrast-aware pre-training framework, ESMSCOP. ESMSCOP leverages a state-of-the-art protein language model to harness evolutionary insights embedded in sequences, and introduces a new encoder to fuse topological and fine-grained spatial structural features. By employing a contrastive pre-training strategy with auxiliary supervision, ESMSCOP effectively bridges the sequence-structure gap, yielding rich and informative representations.

**Conclusions:** Extensive experiments conducted on multiple benchmark datasets demonstrate that ESMSCOP achieves superior performance in protein function prediction tasks compared to existing methods. Furthermore, it shows strong performance even when utilizing relatively less pre-training data compared to some large-scale models.

## 1 Introduction

Proteins are fundamental to virtually all biological processes, including immune response, cellular communication, and molecular transport [1]. Understanding protein functions is thus of great importance for a wide array of downstream applications, such as protein–protein interaction analysis, drug discovery, and precision medicine [2]. A central dogma in molecular biology [3] posits that a protein’s amino acid sequence determines its three-dimensional structure, which in turn dictates its function. Consequently, elucidating the sequence–structure–function relationship of proteins is a fundamental challenge in bioinformatics.

Historically, researchers have inferred protein functions based on sequence similarity, using high-throughput sequencing to discover homologous proteins [4]. However, the scarcity of experimentally validated structure data has hindered the development of structure-based functional inference. With the advent of deep learning, particularly AlphaFold [5, 6] and RoseTTAFold [7, 8], highly accurate 3D protein structures can now be predicted computationally at atomic resolution, significantly reducing the cost and time associated with experimental determination techniques such as X-ray crystallography and cryo-EM [9, 10]. These breakthroughs have galvanized interest in protein representation learning, where machine learning models learn informative embeddings from sequence and structural data to predict protein functions.

The success of representation learning hinges on the choice of protein descriptors [11], which generally fall into two categories: (i) sequence descriptors, which capture the primary structure via amino acid sequences (e.g., FASTA) [12]; and (ii) structure descriptors, which describe the 2D topological or 3D spatial conformations using contact maps, ribbon diagrams, or molecular graphs[13–15]. Consequently, modern protein function prediction methods are typically built upon either sequence-based or structure-based features.

Early sequence-based approaches used handcrafted features to capture evolutionary or physicochemical patterns [16]. Inspired by advances in Natural Language Processing (NLP), deep models such as covolutional neural networks (CNNs) [17], Transformers [12], and BERT-style pretraining [18] have been applied to learn contextualized sequence representations. MSA-based models [19] further improve generalization by incorporating protein family-level information. On the other hand, structure-based methods have leveraged 3D CNNs [20], geometric deep learning [21], and graph neural networks (GNNs)[22–24] to model the intricate spatial and topological properties of protein structures. Some models also utilize surface-level features for more discriminative representations[25].

Recent efforts have explored multi-view learning, aiming to combine complementary perspectives from sequences and structures. For example, Gligorijevic *et al*.[22] integrated GNNs with pre-trained sequence embeddings; Bepler *et al*.[26] incorporated structural signals during sequence model pretraining; Wang *et al*. [27] combined GNNs with language models using structure-guided fine-tuning. These works highlight the importance of learning coherent, structure-aware protein representations. Meanwhile, inspired by self-supervised learning in NLP and computer vision (CV), pretraining has emerged as a dominant strategy in protein modeling. Sequence-based models typically rely on massive corpora of unlabeled data and adopt objectives such as masked language modeling (MLM)[12, 28], contrastive predictive coding (CPC)[29], or next-token prediction [30]. With the increasing availability of structural data, structure-centric pretraining approaches have also flourished, employing geometric contrastive learning [31, 32], bond-level self-prediction [23, 33], or 3D GNN encoders [34].

Despite the impressive progress made in methods for protein function prediction, these methods are still subject to the following limitations:

- Protein annotations are relatively scarce. Annotated data regarding the physicochemical properties and biological functions of proteins are limited [35], as such information typically relies on wet-lab experiments that are time-consuming and costly. This scarcity poses significant challenges for data-driven deep learning approaches. Therefore, it is essential to develop methods that can effectively learn protein representations under unsupervised or weakly supervised conditions.
- Protein structural features are difficult to comprehensively capture. Protein structure plays a fundamental role in determining its function, making accurate structural modeling crucial for function prediction. However, existing sequence-based models [17, 18] often overlook structural information, while most structure-based methods [22] only focus on 2D topological relationships, failing to sufficiently model the geometric conformations in 3D space. As a result, the learned representations may be incomplete and unable to fully capture structure-function relationships. Hence, there is a pressing need for structural modeling approaches that can integrate both topological and spatial features.
- Evolutionary information is challenging to model. Evolutionary information is vital for identifying and inferring protein function, as it highlights conserved regions that are critical for structural stability or biological activity [19]. Multiple sequence alignment (MSA) offers a way to explicitly incorporate evolutionary features, such as conserved residues and co-evolving positions. However, constructing MSAs typically requires a large number of homologous sequences and becomes less effective for low-homology proteins or large-scale datasets due to limited adaptability and computational cost. Existing sequence- or structure-based representations often fail to implicitly capture such evolutionary signals, limiting the model’s ability to generalize functional insights. Thus, there is an urgent need for modeling approaches that can automatically extract evolutionary features from sequences to enhance the generalization and accuracy of function prediction.
- The relationship between protein sequence and structure is underexplored. Protein sequences and structures reflect different levels of biological information: sequences carry evolutionary signals, while structures convey functional mechanisms. However, current approaches typically model proteins from a single perspective or extract sequence and structure features separately before naively concatenating them, without fully leveraging the intrinsic correlation between the two modalities. This lack of cross-modal interaction limits the completeness of the learned representations and may hinder the accuracy of function prediction.

To address these limitations, we propose ESMSCOP, a novel pretraining framework for protein function prediction. ESMSCOP is designed with the following key features:

- We develop a sequence encoder based on ESM-C [36], a state-of-the-art protein foundation model that captures evolutionary information embedded in protein sequences, effectively distilling millions of years of evolution into expressive representations.
- We design a comprehensive protein structure encoder that leverages geometric deep learning to model both topological connectivity and 3D spatial conformations, leading to enriched structural representations with functional relevance.
- We propose a multi-view encoder architecture that jointly learns from both sequence and structure modalities, allowing the model to exploit complementary information across different biological levels.
- We introduce a novel supervised contrastive objective that aligns sequence and structure representations while enforcing consistency across structural conformations, thereby enhancing the robustness and coherence of the learned embeddings.
- We establish a pretraining paradigm that does not rely on explicit functional annotations, enabling effective representation learning under limited label availability and promoting broader applicability to low-resource settings.

A preliminary version of this paper was published in the proceedings of the IEEE International Conference on Bioinformatics and Biomedicine (BIBM) 2024 [32]. Compared to our previous work, we have made several significant improvements in this version. Firstly, we provide a more detailed discussion of the protein function prediction problem, including an enriched review of related work and more rigorous formal definitions. Secondly, we have implemented a novel adaptation of the model architecture by introducing ESM-C, a powerful pre-trained protein language model, to replace the shallow CNN encoder used in the original framework. This change allows for the incorporation of richer evolutionary information into protein representations. Finally, we conducted more comprehensive experimental evaluations. We introduced the new task of protein active site prediction to demonstrate the effectiveness of our proposed updated model, ESMSCOP, across different function prediction tasks. Extensive experiments demonstrate that ESMSCOP consistently outperforms state-of-the-art baselines in both accuracy and robustness across diverse functional prediction tasks.

## 2 Material and methods

### 2.1 Pre-training Datasets

We utilize AlphaFold Swiss-Prot^1^, a large-scale protein structure database predicted by the AlphaFold model [5], for pre-training. This dataset is derived from UniProtKB/Swiss-Prot [37], a manually curated and reviewed protein knowledgebase, and contains approximately 542K high-quality protein 3D structures. Its high structural accuracy and comprehensive coverage make it an ideal resource for self-supervised learning in protein representation tasks.

### 2.2 Benchmark Datasets

#### 2.2.1 Protein Function Prediction

We utilize four benchmark datasets for protein function prediction: Enzyme Commission (EC)[22], Gene Ontology Molecular Function (GO-MF), Gene Ontology Cellular Component (GO-CC), and Gene Ontology Biological Process (GO-BP)[23]. To ensure fair comparison, we follow the dataset splitting protocol proposed by Gligorijević et al. [22], where proteins in the test set share no more than 95% sequence identity with those in the training set. The four datasets are described as follows:

- **EC**: A dataset containing protein Enzyme Commission (EC) numbers, which describe the catalytic activities of enzymes in biochemical reactions.
- **GO-MF**: A dataset annotated with Gene Ontology Molecular Function (MF) terms. MF terms refer to the elemental activities performed by individual gene products (*e*.*g*., binding or catalysis).
- **GO-CC**: A dataset labeled with Gene Ontology Cellular Component (CC) terms, which indicate the locations within the cell where a protein is active (*e*.*g*., mitochondrion, nucleus).
- **GO-BP**: A dataset annotated with Gene Ontology Biological Process (BP) terms. BP terms describe broader biological goals or processes accomplished by multiple molecular activities, such as DNA repair or signal transduction.

#### 2.2.2 Protein Active Site Identification

In this task, models are trained to learn a mapping function that assigns each residue a binary label (0 or 1), aiming to identify potential protein–protein interaction interfaces between the target protein and its binding partners. We adopt the PPBS dataset curated by Tubiana et al. [38], following their data partitioning strategy. Based on the Dockground database of protein–protein interfaces [39], each unique PDB chain involved in one or more interfaces is treated as an individual sample. Chains with fewer than 10 residues or associated with designed proteins are excluded. The validation and test sets in PPBS are split according to the following criteria:

- **70%**: Sequences with at least 70% identity to a sample in the training set.
- **Homo**: Sequences with no more than 70% identity to any training sample, but belonging to the same protein superfamily as at least one training sample.
- **Topo**: Sequences that share similar topological structures (based on the T-level of the CATH classification [40]) with at least one training sample, but are not included in the 70% or Homology sets.
- **None**: Sequences that do not meet any of the above criteria.
- **All**: A combined set including all of the divisions above.

### 2.3 Baselines

To prove the effectiveness of ESMSCOP, 17 approaches are selected as baselines, which can be classified into three categories, namely sequence-based, structure-based, and pretrained models.

- *Sequence-based supervised methods:* ResNet [12] and CNN [17] are two types of the convolution neural network methods. LSTM [12] is one of the recurrent neural networks; Transformer [12] leverages the self-attention mechanism.
- *Structure-based supervised methods:* GCN [41], GAT [42], GVP [43], GraphQA [44], IEConv [31] and GearNet [23] are methods based on graph convolution theory; GearNetEdge-Sup [23] takes the chemical bond features of the protein into account; 3DCNN [20] allows better capture of spatial information.
- *Pre-trained methods:* ProtBERT-BFD [30] and ESM-1b [28] are pre-trained language models; DeepFRI [22] and GearNetEdge-Pre [23] use the structural information of the protein to pre-train the model. LM-GVP [27] utilizes both sequence and structure information to pretrain the model.

### 2.4 Evaluation Metrics

We evaluate ESMSCOP by the area under the precision-recall curve (AUPR) and protein centric maximum F-score (F_max_) [22]. AUPR summarizes the precision-recall curve by calculating the weighted precision at each threshold [45], and it is effective for imbalanced classification. AUPR can be formalized as follows:

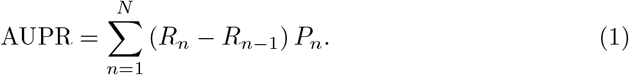

where *R*_*n*_ and *P*_*n*_ are recall and precision of the *n*-th threshold, and *N* is the overall number of thresholds. F_max_ is used to evaluate the accuracy for multi-label classification. In order to calculate F_max_, the precision and recall of each protein are firstly calculated and then averaged across all proteins. Given a target protein *i* and decision threshold *λ* ∈ [0, 1], the precision and recall for this protein is defined as:

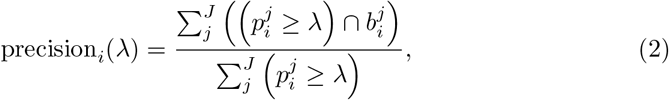

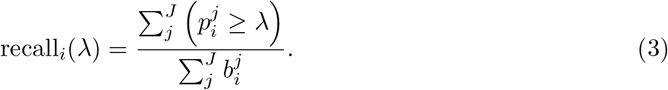

where 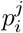 refers to the predicted probability of the *i*-th protein belonging to the *j*-th category, 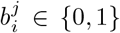 represents the corresponding ground-truth label, and *J* is the total number of categories. The average precision and recall of all proteins can be computed according to the following formula:

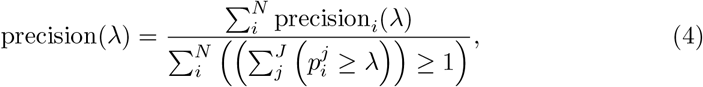

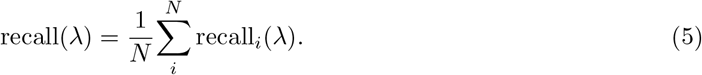

where *N* is the total count of proteins. F_max_ can be defined as:

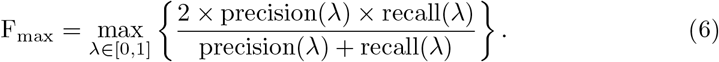

### 2.5 Preliminaries

In this section, we first introduce some fundamental concepts and then describe the task of protein function prediction for better understanding.

#### Definition 1

(Protein Sequence Descriptor) A protein sequence descriptor is a list of symbols that are drawn from a finite set Σ of residue types and arranged in a specific order, denoted as *S* =*< e*_1_, *e*_2_, *e*_3_, *· · ·, e*_*n*_ *>*, where *e*_*i*_ ∈ Σ (1 ≤ *i* ≤ *n*). The sequence of *S* represents the primary structure of a protein.

#### Example 1

Figure 1(a) shows part of the structure of the insulin A-chain represented by a ribbon diagram. The corresponding amino acid sequence is described as GIVEQ in Figure 1(b), where G, I, V, E, Q are the single-letter codes for glycine, isoleucine, valine, glutamic acid, and glutamine, respectively.

**Fig. 1.**
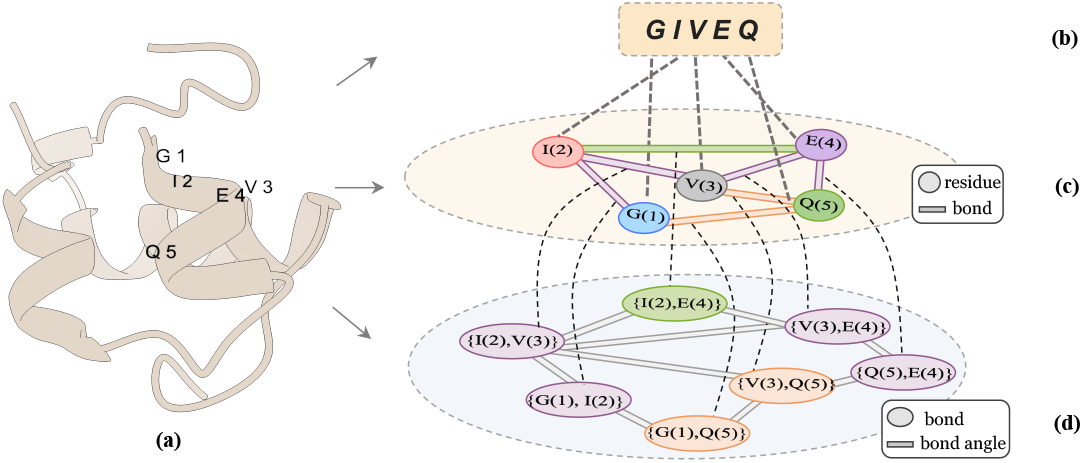
Schematic illustration. (a) Overall structure of insulin, and (b-d) provide annotations for the selected fragment within structure (a). (b) shows its amino acid sequence (GIVEQ), while (c) and (d) depict its corresponding Protein Topological Graph (PTG) and Protein Spatial Graph (PSG), respectively.

#### Definition 2

(Protein Topological Graph) A protein topological graph (PTG) is an undirected graph 𝒢 = (𝒱, ℰ) that represents a protein. Each node *v* ∈ 𝒱corresponds to a residue, and each edge (*u, v*) ∈ ℰ represents a chemical bond between two residues *u* and *v*. We denote the residue attribute of residue *u* as ***x***_*u*_ and the bond attribute of bond (*u, v*) as ***x***_*uv*_. The PTG is a kind of protein structure descriptor.

#### Example 2

Figure 1(c) depicts an instance of the Protein Topological Graph (PTG) for the insulin A-chain fragment shown in Figures 1(a) and (b). In the PTG, each node corresponds to an amino acid residue from this fragment. We can find there are five residues and seven bonds in this graph. Specially, the symbol *G*(1) represents the glycine with index 1 and the symbol {*G*(1), *I*(2)} represents a bond between glycine *G*(1) and isoleucine *I*(2).

#### Definition 3

(Protein Spatial Graph) A protein spatial graph (PSG) is an undirected graph ℳ = (ℰ, 𝒜) that is defined based on the protein topological graph 𝒢. Each node *e*_*uv*_ ∈ *E* in the graph stands for a bond (*u, v*) in the 𝒢, and its attribute is denoted as *x*_*uv*_. Each edge *a*_*uvw*_ ∈ 𝒜 indicates the bond angle between two chemical bonds (*u, v*) and (*v, w*) that share a common residue *v*. It is important to note that the chemical bond angle *a*_*uvw*_ ∈ [0, *π*], and its attribute is denoted as ***x***_*uvw*_. The PSG serves as one of the protein structure descriptors.

#### Example 3

Figure 1(d) shows the PSG of the insulin A-chain. In Figure 1(d), each node stands for a chemical bond that exists in the protein topological graph. As an illustration, the node {*G*(1), *Q*(5)} in Figure 1(d) corresponds to the chemical bond between glycine *G*(1) and glutamine *Q*(5), and the node {*I*(2), *G*(1)} represents the chemical bond between valine *V* (3) and glutamine *Q*(5) in Figure 1(c). The two chemical bonds {*I*(2), *V* (3)} and {*V* (3), *Q*(5)} share one residue, *i*.*e*., valine *V* (3), and the edge *a*_*I*(2)*V* (3)*Q*(5)_ between the two chemical bonds in the Figure 1(d) represents the bond angle between them.

#### Problem 1

(Protein Function Prediction) Consider a set of proteins, denoted as 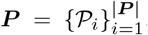, where each protein 𝒫 ∈ ***P*** is a combination of protein sequence descriptors and protein structure descriptors. The labels 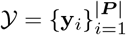of proteins are associated with one or more functional properties. **Protein function prediction** intends to learn the protein representation that can be used to predict its labels by utilizing the diverse information from single or multiple views. The problem is equivalent to finding a function ℱ : ***P*** → 𝒴.

### 2.6 The Design of ESMSCOP

#### 2.6.1 Sequence View Guided Representation

Protein sequences, commonly represented in the FASTA format, serve as the input for our sequence view pathway. We employ the pre-trained ESM-C model, designated as *g*_*seq*_ in Figure 2, to generate sequence-based protein representations. ESM-C’s extensive pre-training on large sequence datasets allows it to capture significant evolutionary context, thereby enhancing the expressiveness of the generated representations.

**Fig. 2.**
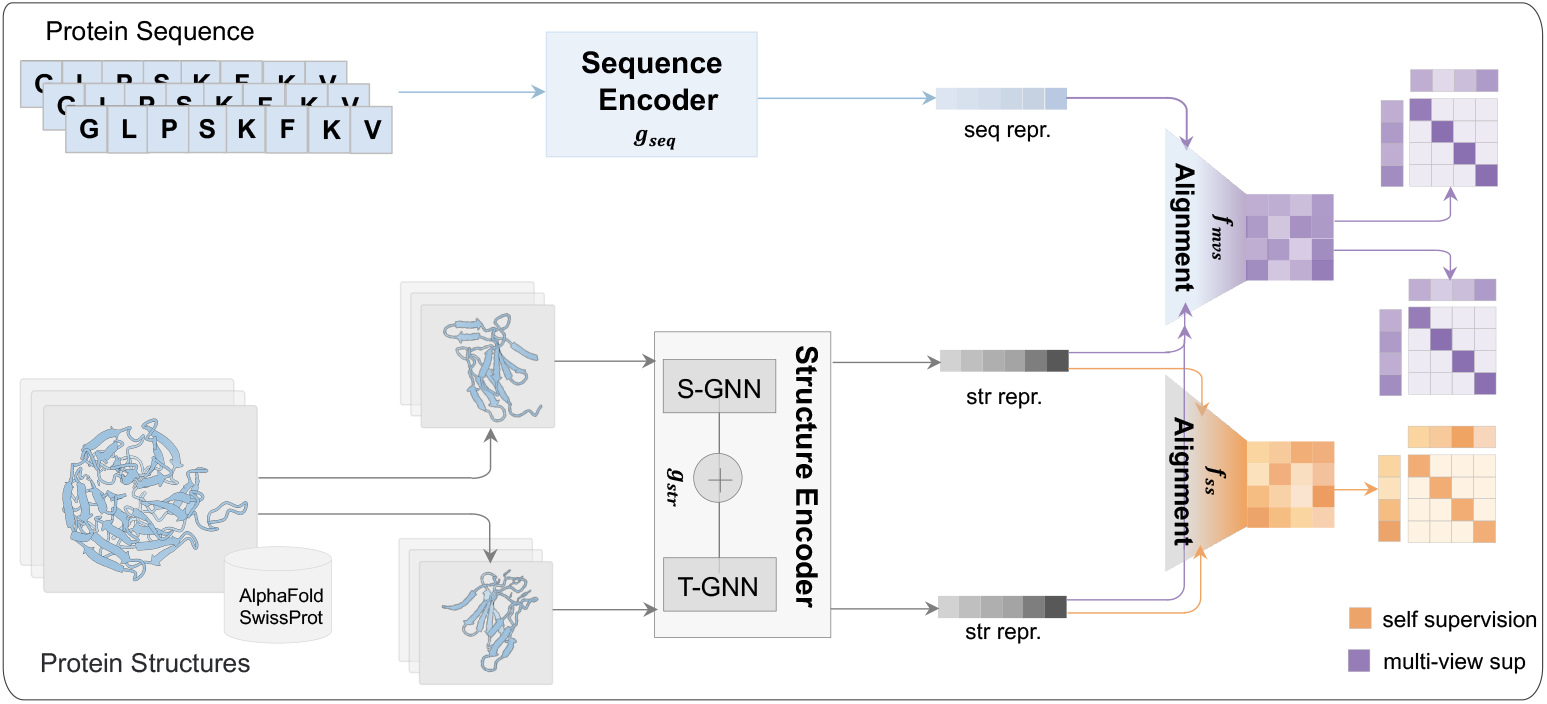
Overview of the proposed ESMSCOP framework. The framework first utilizes a sequence encoder (*g*_*seq*_) to learn sequence representations from protein sequences. Concurrently, protein structures are augmented (or multiple structural views are processed), and a structure encoder (*g*_*str*_, integrating S-GNN and T-GNN) extracts structural representations. These representations are then fed into two alignment modules: a structure-structure alignment module (*f*_*ss*_) aligns representations from different structural views using self-supervision; and a sequence-structure alignment module (*f*_*ms*_) aligns sequence and structure representations using multi-view supervision, projecting them into shared embedding spaces for comparison.

Input FASTA strings *S* are first transformed into dense vectors via an embedding layer *ϕ*(·). The ESM-C model then processes these embeddings to produce the final sequence-view representation ***h***^*S*^:

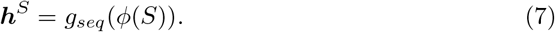

This representation ***h***^*S*^ is subsequently utilized within the pre-training module.

#### 2.6.2 Structure View Guided Representation

For the structural perspective, we utilize Protein Topological Graphs (PTG, denoted as 𝒢) and Protein Spatial Graphs (PSG, denoted as ℳ), as defined in Section 2.5, to capture topological and spatial characteristics, respectively. Chemical bonds, present in both graph types, link these spatial and topological views.

We first employ a Spatial Graph Neural Network (S-GNN) operating on the PSG (ℳ) to compute embeddings ***h***_*uv*_ for each chemical bond. S-GNN iteratively aggregates neighborhood information. Bond embeddings (*u, v*) and (*v, w*) are initialized as 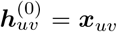 and 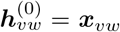. The *k*-th iteration update for bond (*u, v*), denoted 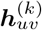, is formalized as:

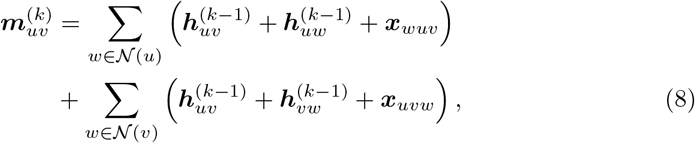

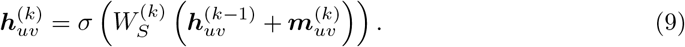

where 𝒩 (*u*) and 𝒩 (*v*) represent the neighbors of *u* and *v*. 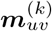 is the intermediate message for bond (*u, v*) at iteration *k*, 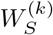 is the *k*-th layer’s learnable matrix, and *σ* is a non-linear activation function (*e*.*g*., ReLU).

Next, a Topological Graph Neural Network (T-GNN) processes the PTG (*G*) to learn residue embeddings. Residue embeddings are initialized as 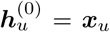. T-GNN iteratively updates the embedding 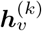 for residue *v* at iteration *k*:

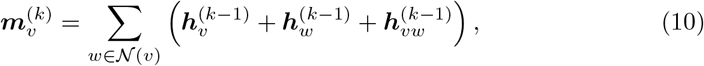

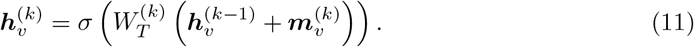

In this update, 𝒩 (*v*) contains neighbors of residue *v*, 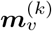 is the intermediate representation for residue *v*, 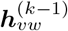 is the (*k* − 1)-th iteration chemical bond embedding from S-GNN (acting on ℳ), and 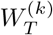 is the *k*-th layer’s learnable weight matrix.

Finally, to obtain a global protein structure representation, we apply average pooling over all residue embeddings after *K* iterations:

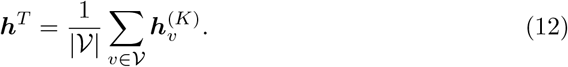

Where *K* is the total number of iterations. The resulting structure-view representation, ***h***^*T*^, generated by the entire structure encoder process denoted as *g*_*str*_ in Figure 2, proceeds to the pre-training module.

#### 2.6.3 Protein Representation Alignment

Sequence and structure representations (***h***^*S*^ and ***h***^*T*^) originate from distinct encoders and inhabit separate embedding spaces, precluding direct comparison. To enable cross-view analysis, we introduce a representation alignment module. This module employs two distinct mapping functions, denoted as *f*_ss_ and *f*_mvs_ in Figure 2, tailored to the specific requirements of self-supervision and multi-view supervision, respectively (detailed in Section 2.6.4). These functions project the initial representations into shared latent spaces.

For instance, the mapping *f*_ss_, used for self-supervision, transforms the structure representation 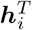 of protein *i* as follows:

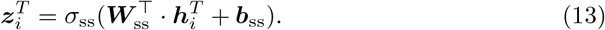

Here, 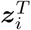 represents the projected embedding. The function *f*_ss_ comprises an activation function *σ*_ss_, a weight matrix ***W***_ss_, and a bias vector ***b***_ss_. An analogous projection yields the transformed sequence representation 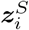.

Similarly, the mapping function *f*_mvs_, designed for multi-view supervision, transforms the structure representation 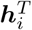 for protein *i* via:

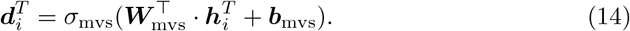

Where 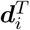 is the resulting projected embedding. The components *σ*_mvs_, ***W***_mvs_, and ***b***_mvs_ parallel those in Equation 13. This function also transforms the sequence representation into 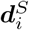 within the same multi-view space.

#### 2.6.4 Auxiliary Supervision for Pre-training

This section details the two auxiliary supervision tasks employed during pre-training: self-supervision within the structure view and multi-view supervision bridging the sequence and structure views.

##### Self-Supervision within Structure View

To capture intrinsic structural properties, we employ a self-supervision strategy based on the NT-Xent loss [46]. For a given protein 𝒫, we generate two distinct structural views, 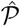 and 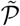, through data augmentation. This involves randomly selecting a residue and defining a sub-protein consisting of all residues within a 10.0 Å sphere centered on it. The self-supervision objective is to maximize the consistency between the representations of these two augmented views derived from the same protein.

Considering a batch of *N* proteins, let 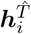 and 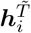 be the structure-view representations of the two sub-proteins from protein *i*, obtained via the structure encoder *g*_*str*_. The mapping function *f*_ss_ projects these into a latent space, yielding 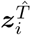 and 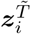. The NT-Xent loss for protein *i* is calculated as:

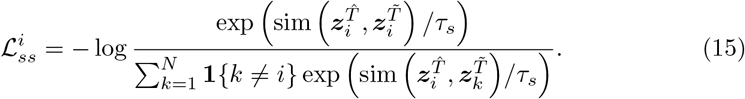

Where *τ*_*s*_ is a temperature hyperparameter, **1** {*k* ≠ *i*} ∈ {0, 1} is an indicator function evaluating to 1 if *k* ≠ *i*, and sim(·,·) denotes the cosine similarity function.

##### Multi-View Supervision within Sequence-Structure View

While self-supervision focuses solely on structural information, multi-view supervision aims to bridge the gap between sequence and structure representations. Since the augmented sub-proteins 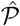 and 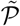 primarily capture local structural features, they may not fully reflect the protein’s global characteristics. Therefore, for this task, we use the representation ***h***^*S*^ derived from the *original* protein sequence *S* (processed by *g*_*seq*_ without augmentation) alongside the sub-structure representations.

For the *i*-th protein in a batch of *N*, we form sequence-structure pairs 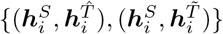. The mapping function *f*_mvs_ projects these representations into the multi-view embedding space, resulting in 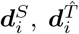 and 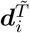. The objective is to maximize the similarity between the sequence representation and its corresponding sub-structure representations. The loss component associated with the sequence encoder for sub-protein 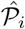 is:

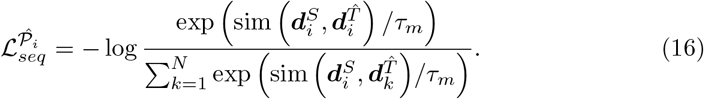

Where *τ*_*m*_ is a temperature hyperparameter. A similar loss, 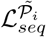 is computed for sub-protein 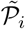. The total sequence encoder loss for protein *i* is their average:

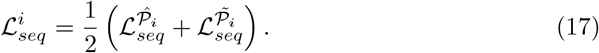

Symmetrically, the loss component for the structure encoder comparing 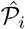 to sequences is:

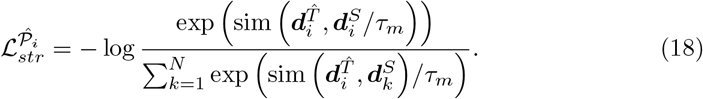

Similarly, we compute 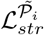 for the other sub-protein. The total structure encoder loss for protein *i* is as follows:

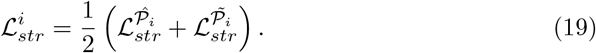

The overall multi-view supervision loss for protein *i* combines these:

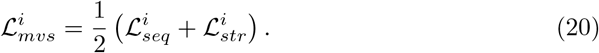

Finally, the total pre-training loss ℒ_*pre*_ combines both supervision signals across the batch:

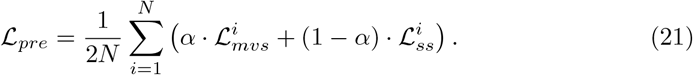

Here, *α* is a hyperparameter balancing the contributions of the multi-view (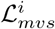) and self-supervision (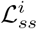) losses. The model (ESMSCOP) is pre-trained by minimizing ℒ_*pre*_ via backpropagation. For downstream tasks like protein function prediction (Section 3), the pre-trained structure encoder *g*_*str*_ is typically used, often augmented with an MLP prediction head.

### 2.7 Implementation Details

We utilize TorchDrug [47] to transform the original PDB file of a protein into the protein topological graph and protein spatial graph. In order to generate initial features for residues, chemical bonds, and bond angles, we partially adopt the approach proposed by Zhang *et al*. [23]. We leverage one-hot encoding to generate the initial residue feature of a protein as a 21-dimensional tensor, representing twenty standard amino acids and one additional unknown amino acid. To represent the initial features of a chemical bond, we concatenate the original features of the two residues which form this bond with a tensor representing the bond length, resulting in a 62-dimensional tensor. The bond length is converted into a 20-dimensional tensor using a radial basis function (RBF). Similarly, the bond angle between two chemical bond is also converted into a 20-dimensional tensor using another RBF. For a protein sequence, each amino acid symbol is embedded as a 16-dimensional dense tensor. These features will be used to train structure-based and sequence-based encoders.

We select the largest batch size that the available GPU memory (24GB) could support, which is 256, and complete the pre-training. The batch size for the benchmark datasets varies depending on the specific dataset. Please refer to Section 3.4 for the best batch size. The temperature coefficient *τ*_*s*_ and *τ*_*m*_ are initialized to 0.07. The hyper-parameter *α* is obtained by a grid search strategy ranging from 0 to 1 with an interval 0.1. The learning rate for pre-training and benchmark datasets is 1e-4, using AdamW optimizer.

Due to the lengthy computational time required for pre-training, we determine the numbers of layers for the structure-based encoder in advance on benchmark datasets, which are set to 6 (see Section 3.4). And it allows us to achieve optimal results within limited computational resources. The sequence-based encoder ESM-C is initialized with pre-trained model weights^2^. We train ESMSCOP from scratch for 50 epochs. The code implementation and datasets are available in the repository^3^.

## 3 Results and discussion

### 3.1 Effectiveness on the Protein Function Prediction

The F_max_ and AUPR scores on the four benchmark datasets regarding the protein function prediction are presented in Table 2, with the best results highlighted in bold and the second-best results underlined. In terms of the F_max_ metric, ESMSCOP consistently outperforms all baseline methods. Specifically, it achieves improvements of 1.5%, 2.9%, and 2.6% and 1.4%, over the respective second-best results across the datasets. Regarding AUPR, ESMSCOP also attains the highest performance on all datasets, with improvements of 2.1%, 2.5%, 1.9% and 1.7%, respectively. These results demonstrate that ESMSCOP benefits significantly from the contrastive pre-training stage, which enhances its ability to capture both the shared patterns and distinctive features of proteins across sequence and structural representations.

**Table 1.** The overview of tasks and and datasets.

| TASK | Dataset | #Train | #Valid | #Test | Metric |
| --- | --- | --- | --- | --- | --- |
| Function Prediction | EC | 15,550 | 1,729 | 1,919 | AUPR+F<sub>max</sub> |
| Function Prediction | GO-MF | 29,899 | 3,322 | 3,415 | AUPR+F<sub>max</sub> |
| Function Prediction | GO-CC | 29,899 | 3,322 | 3,415 | AUPR+F<sub>max</sub> |
| Function Prediction | GO-BP | 29,899 | 3,322 | 3,415 | AUPR+F<sub>max</sub> |
| Binding Site Prediction | PPBS | 12,577 | 3,178 | 3,984 | AUPR |

**Table 2.** The performance comparison with baselines on benchmark datasets under metrics F_max_ and AUPR about protein function prediction task. [*] denotes results taken from [27].

| | Method | Pre-training Dataset<br>/# Parameters | $F_{\max}$ | | | | AUPR | | | |
| --- | --- | --- | --- | --- | --- | --- | --- | --- | --- | --- |
|  |  |  | EC | GO-BP | GO-MF | GO-CC | EC | GO-BP | GO-MF | GO-CC |
| Sequence | CNN [17] | - / 9M | 0.545 | 0.244 | 0.354 | 0.287 | 0.526 | 0.159 | 0.351 | 0.204 |
|  | ResNet [12] | - / 11M | 0.605 | 0.280 | 0.405 | 0.304 | 0.590 | 0.205 | 0.434 | 0.214 |
|  | LSTM [12] | - / 27M | 0.425 | 0.225 | 0.321 | 0.283 | 0.414 | 0.156 | 0.334 | 0.192 |
|  | Transformer [12] | - / 21M | 0.238 | 0.264 | 0.211 | 0.405 | 0.218 | 0.156 | 0.177 | 0.210 |
| Structure | GCN [41] | - / 1M | 0.320 | 0.252 | 0.195 | 0.329 | 0.319 | 0.136 | 0.147 | 0.175 |
|  | GAT [42] | - / 3M | 0.368 | 0.284* | 0.317* | 0.385* | 0.320 | 0.171 | 0.329 | 0.249 |
|  | GVP [43] | - / 6M | 0.489 | 0.326* | 0.426* | 0.420* | 0.482 | 0.224 | 0.458 | 0.279 |
|  | 3DCNN [20] | - / 5M | 0.077 | 0.240 | 0.147 | 0.305 | 0.029 | 0.132 | 0.075 | 0.144 |
|  | GraphQA [44] | - / 2M | 0.509 | 0.308 | 0.329 | 0.413 | 0.543 | 0.199 | 0.347 | 0.265 |
|  | IEConv [31] | - / 5M | 0.735 | 0.374 | 0.544 | 0.444 | 0.775 | 0.273 | 0.572 | 0.316 |
|  | GearNet [23] | - / 11M | 0.730 | 0.356 | 0.503 | 0.414 | 0.751 | 0.211 | 0.490 | 0.276 |
|  | GearNetEdge-Sup [23] | - / 21M | 0.810 | 0.403 | 0.580 | 0.450 | 0.872 | 0.251 | 0.570 | 0.303 |
| Pretrained | DeepFRI [22] | Pfam (10M) / 709K | 0.631 | 0.399 | 0.465 | 0.460 | 0.547 | 0.282 | 0.462 | 0.363 |
|  | ESM-1b [28] | UniRef50 (24M) / 650M | 0.862 | 0.452 | 0.657 | 0.477 | 0.889 | 0.342 | 0.639 | 0.384 |
|  | ProtBERT-BFD [30] | BFD (2.1B) / 420M | 0.838 | 0.279* | 0.456* | 0.408* | 0.859 | 0.188 | 0.464 | 0.234 |
|  | LM-GVP [27] | UniRef100 (216M) / 424M | 0.664 | 0.417* | 0.545* | 0.527* | 0.710 | 0.302 | 0.580 | 0.423 |
|  | GearNetEdge-Pre [23] | Swiss-Prot (542K) / 21M | <u>0.878</u> | <u>0.462</u> | 0.639 | 0.465 | 0.885 | 0.283 | 0.596 | 0.336 |
|  | **ESMSCOP** | Swiss-Prot (542K) / 26M | **0.893** | **0.491** | **0.683** | **0.541** | **0.910** | **0.367** | **0.658** | **0.440** |

Compared with both sequence-based and structure-based models, ESMSCOP consistently outperforms all baselines, demonstrating that the pre-training phase significantly enhances its performance, particularly on small-scale datasets. We further assess the effectiveness of ESMSCOP against other pre-trained models. According to the F_max_ and AUPR metric, ESMSCOP achieves the best performance across all four benchmark datasets.

ESMSCOP consistently outperforms nearly all existing pre-trained models. We attribute this performance gain to its ability to integrate multimodal information from both protein sequences and structures. Specifically, the sequence encoder leverages the pre-trained weights of ESM-C, which captures rich evolutionary signals learned from millions of protein sequences across evolutionary timescales. In addition, the proposed multi-view pre-training objective allows the model to uncover intrinsic correlations between structural and sequential representations, providing informative self-supervised signals that further enhance representation learning. Finally, by incorporating the AlphaFold Swiss-Prot structure dataset, we demonstrate that the evolutionary knowledge embedded in the ESM-C model can be effectively transferred and fused into the structure-based encoder, enabling seamless cross-modal information integration.

Meanwhile, we also provide the number of parameters for each pretrained model in Table 2. For deep learning models, having more parameters usually means higher computational demands and stronger learning capabilities. As indicated in the Table 2, pretrained models generally have more parameters and better performance compared to models based solely on sequence or structure, which highlights the advantage of pretrained models in learning protein representations. Thanks to the distillation of evolutionary information from the sequence-based encoder into the structure-based encoder during pre-training, ESMSCOP achieves strong performance with a relatively small number of parameters. This highlights its advantage in delivering high accuracy while maintaining lower computational and memory costs.

### 3.2 Effectiveness on Active Sites Identification in Protein-protein Interaction

To further evaluate the effectiveness of ESMSCOP, we conducted experiments on the PPBS dataset for protein active site prediction. This task aims to identify key residues involved in protein-protein interactions, which is crucial for understanding protein function. All experiments were conducted under the same baseline settings, and the results are summarized in Table 3. Across all five benchmark subsets, ESMSCOP consistently achieves the best performance.

**Table 3.** The performance comparison with baselines on benchmark datasets under AUPR regarding protein active site prediction.

|  | Method | AUPR |  |  |  |  |
| --- | --- | --- | --- | --- | --- | --- |
|  |  | PPBS-70 | PPBS-Homo | PPBS-Topo | PPBS-None | PPBS-All |
| Sequence | CNN [17] | 0.551 | 0.533 | 0.529 | 0.513 | 0.532 |
|  | ResNet [12] | 0.577 | 0.550 | 0.558 | 0.539 | 0.556 |
|  | LSTM [12] | 0.574 | 0.542 | 0.536 | 0.520 | 0.543 |
|  | Transformer [12] | 0.580 | 0.557 | 0.549 | 0.531 | 0.554 |
| Structure | GCN [41] | 0.684 | 0.635 | 0.620 | 0.547 | 0.622 |
|  | GAT [42] | 0.675 | 0.657 | 0.630 | 0.539 | 0.625 |
|  | GVP [43] | 0.684 | 0.665 | 0.643 | 0.541 | 0.633 |
|  | 3DCNN [20] | 0.624 | 0.558 | 0.541 | 0.509 | 0.558 |
|  | GraphQA [44] | 0.654 | 0.638 | 0.627 | 0.581 | 0.625 |
|  | IEConv [31] | 0.703 | 0.695 | 0.660 | 0.597 | 0.664 |
|  | GearNet [23] | 0.695 | 0.679 | 0.637 | 0.549 | 0.640 |
|  | GearNetEdge-Sup [23] | 0.697 | 0.688 | 0.656 | 0.560 | 0.650 |
| Pretrained | DeepFRI [22] | 0.676 | 0.702 | 0.685 | 0.613 | 0.669 |
|  | ESM-1b [28] | 0.689 | 0.662 | 0.690 | 0.604 | 0.661 |
|  | ProtBERT-BFD [30] | <u>0.725</u> | 0.701 | 0.694 | 0.624 | 0.686 |
|  | LM-GVP [27] | 0.704 | 0.693 | 0.674 | 0.630 | 0.675 |
|  | GearNetEdge-Pre [23] | 0.722 | <u>0.718</u> | <u>0.709</u> | <u>0.633</u> | <u>0.696</u> |
|  | **ESMSCOP** | **0.757** | **0.729** | **0.734** | **0.653** | **0.718** |

Bold and underline denote the best and second-best results, respectively.

This superior performance can be attributed to the model’s ability to capture the topological and spatial structural features of proteins. Since active site prediction relies heavily on accurate modeling of 3D structural characteristics, ESMSCOP’s capability to jointly learn from sequence and structure views enables more precise localization of functional residues.

To provide an intuitive understanding of the model’s predictions, we visualize two randomly selected protein complexes from the test set. For each complex, the activity of amino acid residues is predicted and color-coded, with low activity in gray and high activity in blue. As shown in Figure 3, the predicted active regions closely align with the true interaction interfaces, clearly highlighting potential active sites. These visualizations offer valuable insight into protein functions such as catalysis and signal transduction, further demonstrating the practical utility of ESMSCOP.

**Fig. 3.**
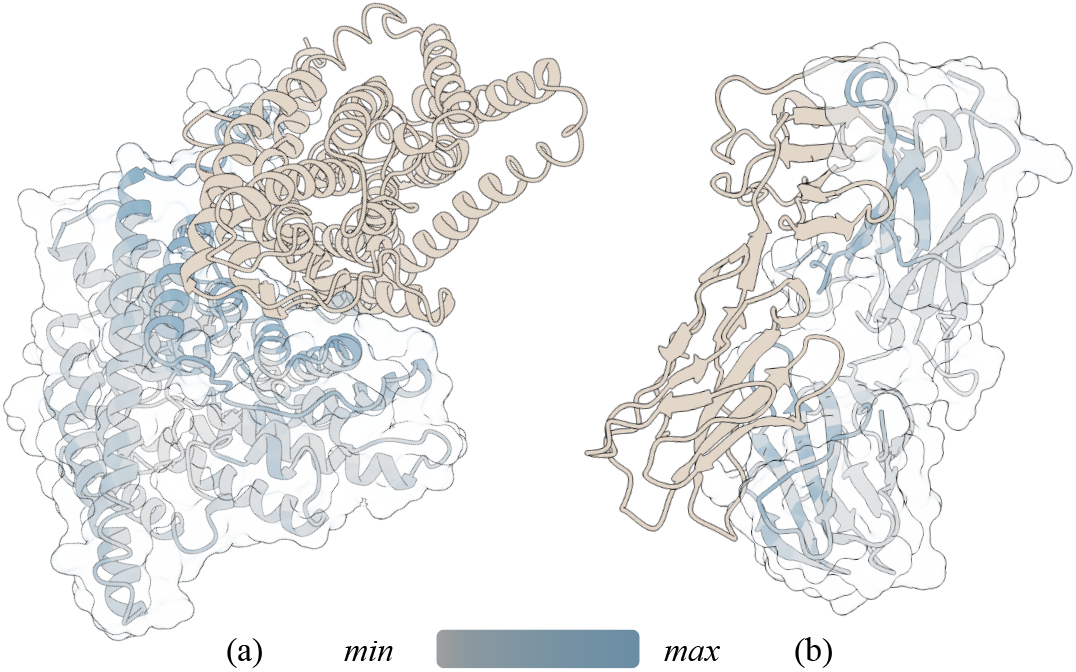
Case study of protein active site prediction. Protein complexes (a) 1KPK and (b) 3DXA are randomly selected from the test set. ESMSCOP predicts the activity of each amino acid residue, with low activity shown in gray and high activity in blue. The predicted active sites align well with the actual binding interfaces, demonstrating the model’s ability to capture functionally relevant regions.

### 3.3 Ablation Study

To gain a deeper understanding of ESMSCOP and validate its effectiveness, we propose three variants of the model: (1) **NSS** (<u>N</u>o <u>S</u>elf-<u>S</u>upervision): ESMSCOP pretrained without self-supervision; (2) **NMV** (<u>N</u>o <u>M</u>ulti-<u>S</u>upervision): ESMSCOP pretrained without multi-view supervision; (3) **NSP** (<u>N</u>o protein <u>SP</u>atial information): ESMSCOP pretrained without protein spatial information.

Figure 4 illustrates the performance of ESMSCOP and its variants on four benchmark datasets. We can see that the F_max_ of ESMSCOP is the highest on these datasets, proving the protein spatial information and pre-training supervision tasks developed in this paper are effective.

**Fig. 4.**
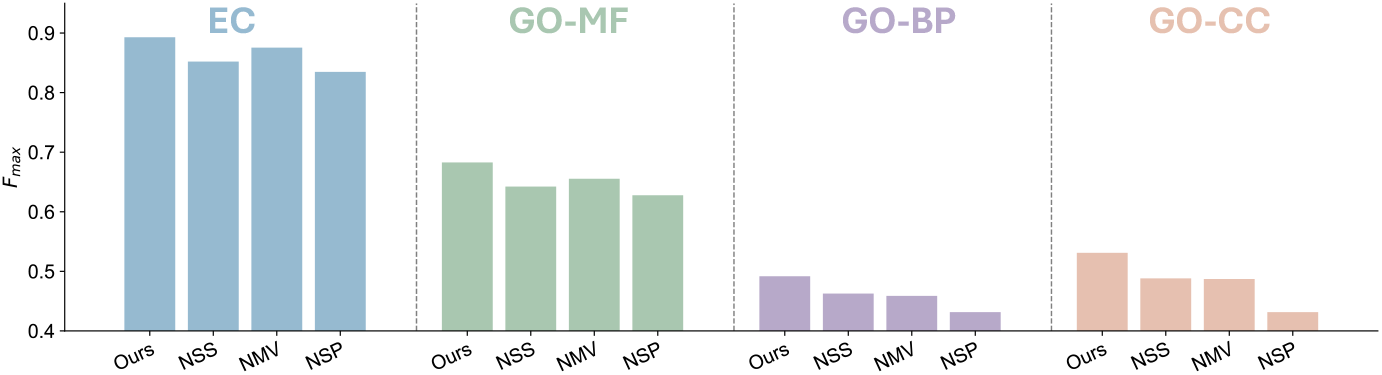
Ablation study of the proposed model. The performance of the model on four protein function prediction datasets is evaluated after removing key components, including self-supervision, multiview supervision, and structural information learning. The results highlight the contribution of each module to the overall performance.

According to the experimental results, it is apparent that the model without spatial information (NSP) has the worst performance among all the model variants, indicating the importance of spatial information for protein function prediction.

Meanwhile, the contributions of self-supervision (SS) and multi-view supervision (MVS) are different on the four benchmark tasks. The SS provides more improvement compared to MVS in GO-MF datasets, but the improvement is close in GO-BP and GO-CC datasets. This is probably due to the nature of different datasets, for example, the EC dataset is used to predict the catalysis of biochemical reactions of proteins, which focuses more on the local structure features of proteins, and SS is able to provide more information about the local structure of proteins compared to MVS. In conclusion, each of the three variants for ESMSCOP plays an essential role in protein function prediction, resulting in a substantial improvement in the experimental results.

### 3.4 Parameter Sensitivity Analysis

Since ESMSCOP primarily relies on the structure-based encoder, we first determine the optimal number of layers *l*_*str*_ for this encoder on downstream tasks to reduce the computational cost of pre-training. Additionally, the pre-training loss function is controlled by a balancing coefficient *α*, which needs to be carefully selected. Furthermore, we investigate the appropriate batch size *b* for the structure-based encoder. To ensure fairness, all other parameters are kept fixed during each individual experiment.

We begin by analyzing the effect of the number of layers in the structure-based encoder. The model achieves consistently strong performance across all tasks when *l*_*str*_ ≥5. While increasing the number of layers can improve model capacity, it does not always result in significant gains and may lead to increased computational complexity, reduced efficiency, and a higher risk of overfitting.

Next, we analyze the sensitivity of the model to the balancing coefficient *α* in the pre-training loss, as shown in Figure 5. The optimal values of *α* are found to be 0.4, 0.2, 0.4, and 0.8 on the EC, GO-MF, GO-CC, and GO-BP datasets, respectively. These differences may stem from the varying characteristics of the datasets and the extent to which the target functions rely on sequence or structure representations. Overall, the performance of ESMSCOP remains relatively stable across a wide range of *α* values. Lastly, we investigate the impact of batch size *b* for the structure-based encoder.

**Fig. 5.**
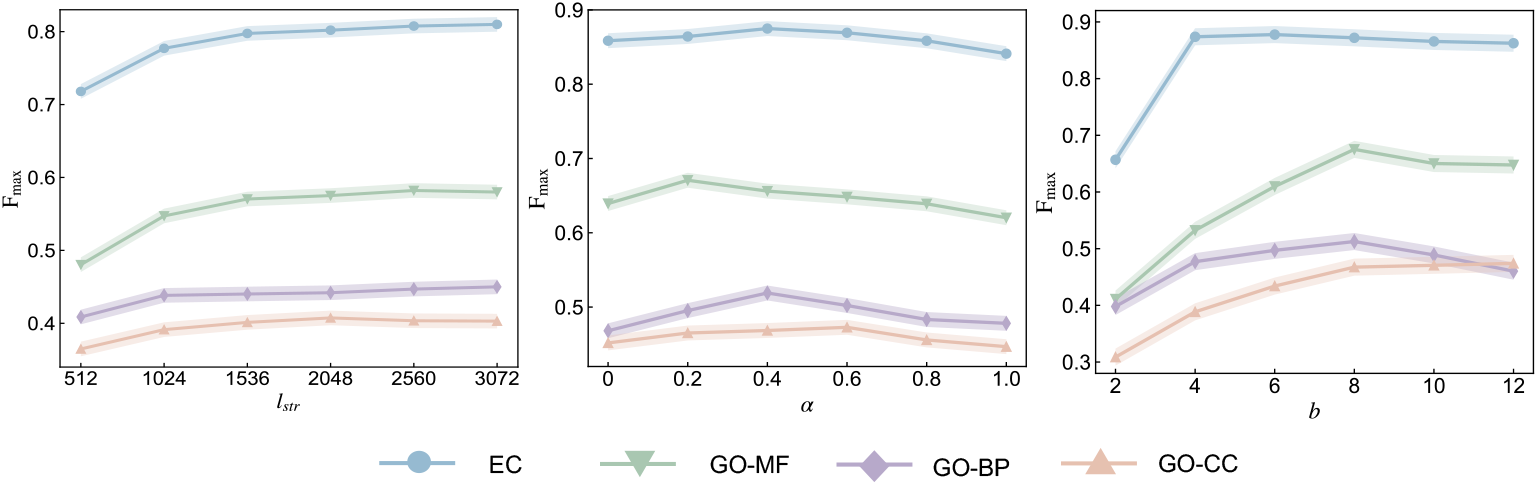
The impacts of the hyperparameters.

The results show that model performance generally improves with larger batch sizes. Specifically, the EC dataset achieves the best performance with a batch size of 6, GO-MF and GO-CC perform best at a batch size of 8, and GO-BP achieves optimal results at a batch size of 12. In general, setting the batch size to 8 or higher yields consistently satisfactory performance across tasks.

## 4 Case Study

Glycoproteins are ubiquitous across various organisms and play critical roles in biological processes such as signal transduction and cell recognition [48]. Accurate identification of glycoproteins is essential for advancing our understanding of disease mechanisms and progression. To this end, we construct a glycoprotein dataset and evaluate the effectiveness of our proposed model, ESMSCOP, for glycoprotein prediction.

The dataset is curated from the RCSB PDB [49], comprising approximately 600 glycoproteins as positive samples and 900 proteins confirmed not to be glycoproteins (lacking glycosylation) as negative samples. We evaluate several models on this dataset, including GearNetEdge-Pre, ESM-1b, and three variants of our method: ESMSCOP-S, ESMSCOP-T, and ESMSCOP. These variants correspond to pretraining on different views—sequence only, structure only, and a joint sequence-structure view, respectively.

To further investigate the quality of the learned representations, we visualize the embeddings using t-SNE [50], a popular technique for projecting high-dimensional data into two-dimensional space. As shown in Figure 6, protein samples are visualized according to their oligosaccharide-binding capability. To quantitatively assess the quality of clustering, we compute the Davies-Bouldin index [51], where a lower score indicates better separation between clusters. The scores for the different methods are displayed in Figure 6. Among all models, ESMSCOP achieves the lowest Davies-Bouldin score (4.0436), outperforming ESMSCOP-T (5.1873), ESM-1b (5.5261), GearNetEdge-Pre (5.5261), and ESMSCOP-S (4.9915). This demonstrates the advantage of integrating both sequence and structural information, enabling ESMSCOP to learn more discriminative and biologically meaningful representations.

**Fig. 6.**
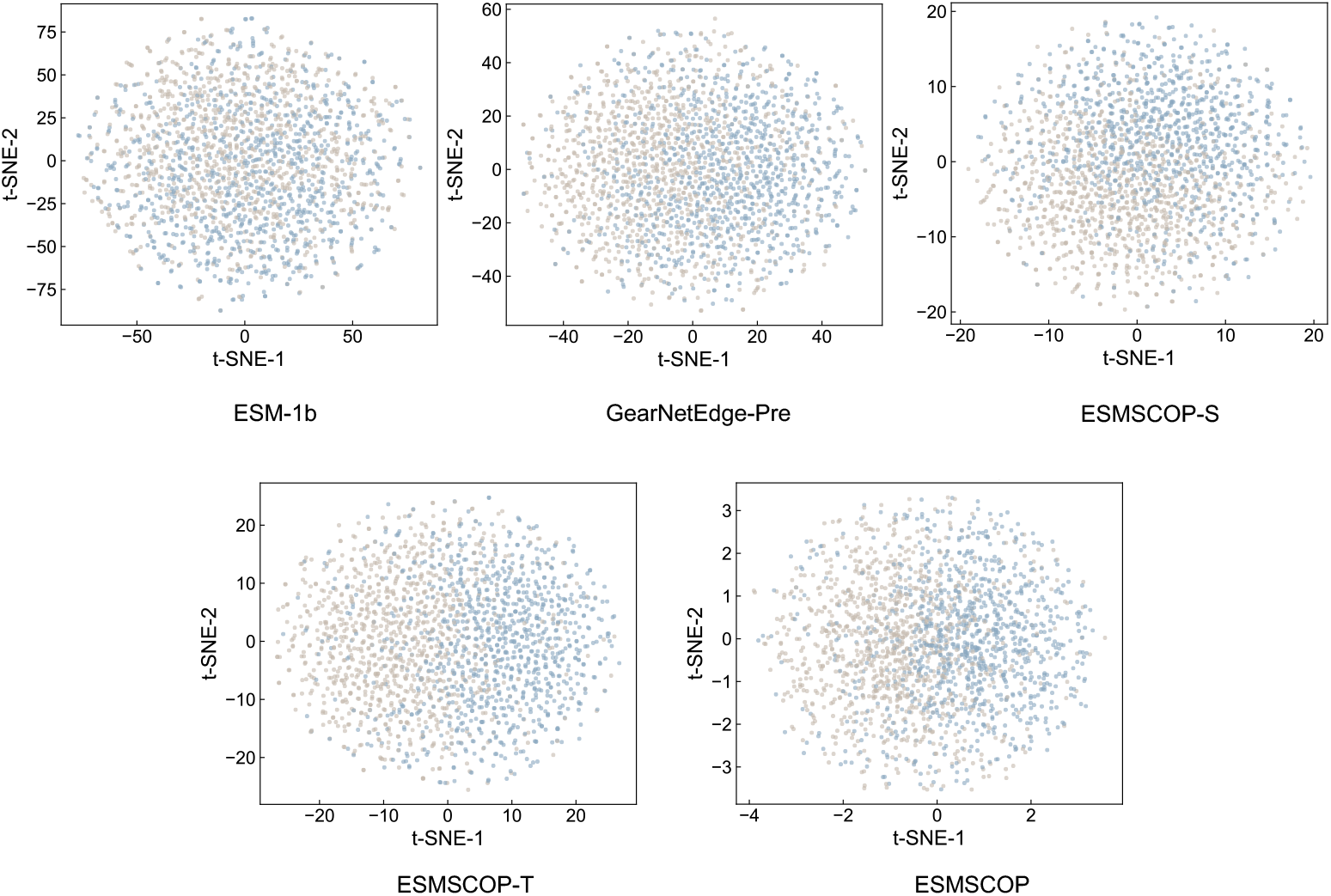
t-SNE visualization comparing protein embeddings from different methods on the glycoprotein dataset. The methods shown are ESM-1b, GearNetEdge-Pre, ESMSCOP-S, ESMSCOP-T, and ESMSCOP, with corresponding Davies-Bouldin index scores of 5.0164, 5.5261, 4.9915, 5.1873, and 4.0436, respectively. Lower scores indicate better clustering separation.

Moreover, as visualized in Figure 7, we observe that ESMSCOP tends to group proteins belonging to the same family or superfamily and clearly separates proteins from different families. For instance, distinct protein families—such as the EGFR family, GPCR family, Protease family, and the Globin family—are well separated in the t-SNE plot generated by ESMSCOP. Since proteins within the same family often share similar biological functions, these observations further validate that the representations learned by ESMSCOP are aligned with real-world biological semantics.

**Fig. 7.**
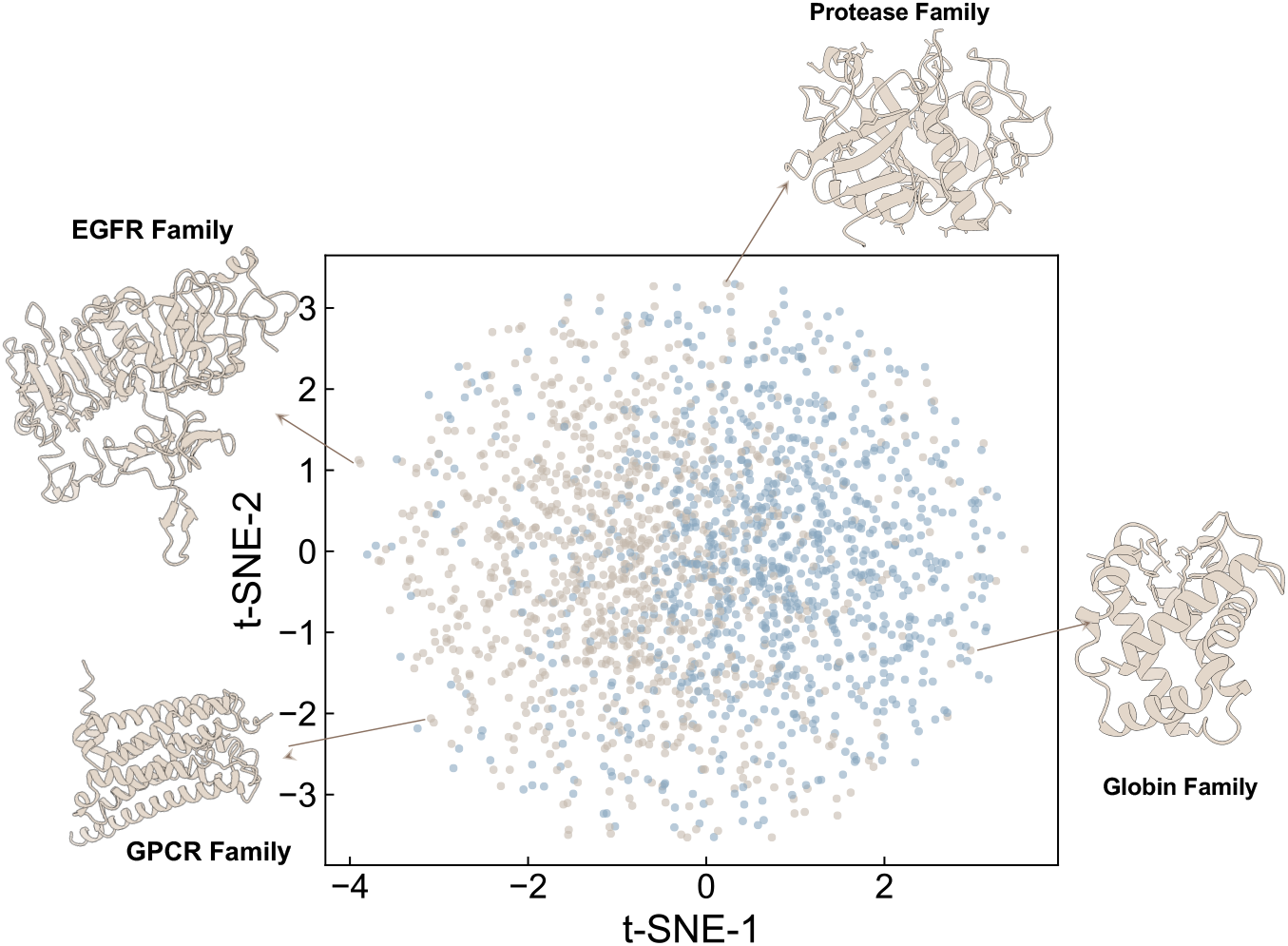
Protein family classification visualized using t-SNE embeddings from ESMSCOP. Proteins from the same family or superfamily (e.g., the illustrated GPCR, EGFR, Protease, Globin families) tend to form distinct clusters, indicating the biological relevance of the learned representations.

## 5 Conclusions

In this work, we propose ESMSCOP, a contrastive-supervised pretraining framework for protein function prediction. We introduce a novel protein encoder that captures both topological and spatial information from the structural view of proteins. Leveraging the latest protein foundation model ESM-C, our framework incorporates both self-supervision and multi-view supervision during pretraining, effectively distilling evolutionary information accumulated over millions of years into the structure-based encoder. This enables the interaction and fusion of sequence and structure representations, revealing cross-modal correlations and facilitating the learning of more comprehensive protein embeddings. Extensive experiments demonstrate that ESMSCOP consistently outperforms existing methods, even when pretrained on smaller datasets.

As to future work, we aim to improve the structure-based encoder by employing E(3)-equivariant models, which will capture the structural features of proteins more effectively and yield highly accurate protein representations. Moreover, we intend to further explore the relationship between pre-trained protein language models and structural models to boost our model’s performance. We believe this research will serve as an inspiration for researchers and expedite the development of new drugs.

## Declarations

### Ethics approval and consent to participate

Not applicable.

### Consent for publication

Not applicable.

### Availability of data and materials

The source code of ESMSCOP and the data related to experiments are stored in a public GitHub repository: https://github.com/mrzzmrzz/ESMSCOP. The open-source weights of ESM-C can be downloaded via this link: https://huggingface.co/EvolutionaryScale/esmc-300m-2024-12.

### Authors’ contribution

R.M. and C.H. wrote the manuscript. R.M. designed the entire experiment and performed the experimental validation. Z.Z. assisted with drawing the illustrations. L.D. supervised the entire project. All authors participated in the revision of the paper.

## Acknowledgements

This work has received support from the School of Computer Science at Sichuan University, for which we are grateful.

## Footnotes

1 https://www.alphafold.com/download

2 https://huggingface.co/EvolutionaryScale/esmc-300m-2024-12

3 https://github.com/mrzzmrzz/ESMSCOP

